# A mitochondria-targeted PPR protein restores cytoplasmic male sterility by post-transcriptional modification of *ORF147* in *Cajanus cajanifolius*

**DOI:** 10.1101/2024.05.17.594745

**Authors:** Joorie Bhattacharya, Rahul B Nitnavare, Richa K. Yeshvekar, Dumbala Srinivas Reddy, Vidhi Sapara, Yogendra Ramtirtha, Yogendra Kalenhalli, Pradeep Reddy, Pooja Bhatnagar-Mathur, Palakolanu Sudhakar Reddy

## Abstract

Restoration factors (Rfs) belonging to the pentatricopeptide repeat proteins (PPRs) family play an essential role in plant growth and development including their binding to CMS-associated mitochondrial RNAs leading to fertility restoration. The present study identified 22 mitochondrial-specific PPRs in pigeonpea and explored the underlying mechanisms of restoration of fertility in the A_4_ CMS system through yeast-three hybrid studies. The identified gene was functionally validated through transgenic expression in *Arabidopsis* model system and obtained conclusive evidence that the identified *Rf-PPR* was responsible for fertility restoration. The sub-cellular localization studies implied that the identified *Rf-PPR* is mitochondrial targeting. The study demonstrated that due to the interaction between mitochondrial CMS mRNA and nuclear Rf-PPR protein, post-transcriptional modification occurred, leading to the inability to translate and accumulate cytotoxic CMS protein resulting in fertility restoration. The study specifically looks into the RNA-protein interaction occurring at the nucleo-cytoplasmic level in the A4 cytoplasm of *Cajanus cajanifolius*.

**Highlights:** The study identifies the restoration of fertility genes corresponding to the CMS-causing *orf147* gene.

## 1. Introduction

Pigeonpea is an important legume produced in a land area of 6.35 million hectares with a global annual production of 5.47 million tons (FAO, 2021). It is an important crop in the tropical and sub-tropical regions of Asia, Africa and South America (Saxena *et al*., 2017). India is the leading producer of pigeonpea (4.32 million tons), contributing to approximately 78.87% of global production (FAO, 2021). Despite its high demand and nutritional value, pigeonpea is categorized as an underutilized crop, possibly due to its geographical limitation (Fatokimi and Tanimonure, 2021). Additionally, while traits such as resistance to disease and earliness have been exploited for crop improvement in pigeonpea, yield remains a matter of concern (Saxena *et al*., 2017).

Pigeonpea is a naturally cross-pollinating legume and exhibits large genetic variation (20-70%), posing significant challenges in maintaining the genetic purity of genotypes. However, this out-crossing characteristic has been used in hybrid breeding technology through male sterile line development for generating high-yielding pigeonpea populations (Dalvi *et al*., 2010). In pigeonpea, three types of male sterility systems exist: genetic (GMS), cytoplasmic-nuclear (CMS) and temperature-sensitive (TGMS) male sterility systems. The CMS system has been investigated thoroughly, and A_4_ cytoplasm has been utilized to generate commercial hybrids (Saxena *et al*., 2005, 2013).

As a maternally inherited trait, cytoplasmic male sterility affects pollen viability and functional male reproductive organs due to dysfunctional mitochondrial chimeric open reading frames (*orf*) (Chase, 2007; Chen *et al*., 2017). These orfs often co-transcribe with other mitochondrial genes, contributing to their genome stability. The restoration of male fertility has been known to occur due to certain nuclear factors called restorer of fertility (*Rf*) genes targeted mitochondrially, leading to a reduction in male fertility-causing CMS proteins (Dahan and Mireau, 2013). Most Rf genes identified in different crop species are from a class of RNA-binding proteins known as the pentatricopeptide repeat proteins (PPR) family. Mitochondrial targeting and binding to the CMS-associated mitochondrial RNA, reduces their accumulation. (Bentolila, 2002; Brown *et al.,* 2003; Desloire *et al.,* 2003; Koizuka *et al.,* 2003; Klein *et al.,* 2005; Wang *et al.,* 2006; Hu *et al.,* 2012). PPR binding also influenced RNA secondary and tertiary structures to make hidden sights for binding more accessible. All in all, the CMS-Rf interaction mechanism has served as a model to study the nucleo-mitochondrial interactions in crop systems (Dahan and Mireau, 2013)

Our previous study identified a novel mitochondrial *orf147,* responsible for CMS in pigeonpea, and caused aberrant floral development, dysfunctional anther dehiscence, reduced lignin content in anthers and programmed cell death (PCD) (Bhatnagar-Mathur *et al*., 2018). This 444 nucleotide pigeonpea ORF also caused partial sterility in transgenic chickpea (*Cicer arietinum*) expressing *orf147* (Bhattacharya *et al*., 2023). The identification of *orf147*-causing CMS in pigeonpea paved the way for exploring the underlying mechanisms of restoration of fertility corresponding to *orf147*. This study attempts to determine *Rf-PPR* corresponding to *orf147, which is* responsible for the restoration of fertility in pigeonpea.

## Materials and methods

### Identification of mitochondrial targeting PPRs in *Cajanus cajanifolius*

The pentatricopeptide repeat genes (PPR) genes were retrieved from the Pigeonpea (*Cajanus cajanifolius)* PPR genes database (https://ppr.plantenergy.uwa.edu.au/ppr/show_species/Cajanus%20cajan) and crosschecked with NCBI database by using BLAST search approach (https://blast.ncbi.nlm.nih.gov/Blast.cgi). The overlapping and duplicate genes were removed. The *in-silico* sub-cellular localisation of the PPRs was done and the ones targeting the mitochondria were shortlisted using tools such as Mitoprot (https://ihg.gsf.de/ihg/mitoprot.html). In addition, the organelle targeting domains of PPR proteins were predicted using TargetP1.1server (http://www.cbs.dtu.dk/services/TargetP/) Predotar (https://urgi.versailles.inra.fr/predotar/). In cases of indistinctness between the results of the three predicting softwares, the prediction with the better confidence was retained and used for expression analysis. The chromosome mapping of mitochondrial PPR genes was done by using information available on public databases like National Centre for Biotechnology Information (https://www.ncbi.nlm.nih.gov/), NCBI and generated using MapChart-WUR (https://www.wur.nl/en/show/mapchart.htm).

### Plant material

The seeds of *Cajanus cajanifolius* male sterile line (A; ICPA2039), its restorer (R; ICPL 87119), two short- and medium-duration restorer lines (ICPL 161 and ICPL 20098, respectively) and a non-restorer line (ICPL 87091) were obtained from pigeonpea breeding unit at International Crops Research Institute for the Semi-Arid Tropics (ICRISAT), Hyderabad, India.

### Expression profiling of putative Rfs

The seeds were grown in a controlled environment, and total RNA was isolated from young, unopened flower buds (100mg) using RNeasy Plant Mini Kit (Qiagen, Germany), followed by cDNA synthesis using Thermoscript RT-PCR system (Invitrogen, USA) Kit. The expression profiles of shortlisted PPRs were analysed in sterile A-line and restorer R-lines using the Realplex Real-Time PCR system (Eppendorf, Germany) and SYBR Green mix (Bioline) in 96 well optical reaction plates (Axygen, USA) sealed with ultra-clear sealing film (Platemax) for 40 cycles using gene-specific primers (Supplementary Table S1) and normalized using *PTB1*and *IF3* reference primers (Supplementary Table S1 and S2) (Sinha *et al.,* 2015). The relative expression was calculated through the 2^-ΔΔ^CT method (Winer *et al*., 1999).

### Yeast three-hybrid (Y3H) studies

The shortlisted putative Rfs identified through expression profile analyses were further taken for three yeast hybrid studies to identify interacting partners with *orf147*. Y3H enables the study of RNA and protein interaction. Briefly, interaction studies of *Rf* and *orf147* were carried out using components of Matchmaker Gold Yeast-two hybrid system, Takara using MS2 protein expressing in bait vector (pGBKT7), MS2-orf147 hybrid RNA expressing in pPRP1 vector and identified PPRs expressing in prey vector (pGADT7) All three components were then co-transformed into yeast host (*Saccharomyces cerevisiae* strain Y187 (Takara) at equimolar concentrations in DDO/X/A [SD/–Leu/–Trp/X-a-Gal/AbA] media (containing kanamycin to prevent bacterial contamination) through electroporation and incubated at 30°C for approximately 3-4 days until appearance of blue colonies. AD was used as a positive control, an empty vector as a negative control, and independent colonies counted as positive interactions (Hamid and Ismail, 2018). After the three-hybrid screening using yeast mating, plasmids of randomly picked blue colonies were rescued using Easy Yeast Plasmid Isolation Kit, Takara and sent for sequencing to confirm positive colonies. The positive *Rfs* determined through sequencing were then taken further for functional validation.

### Determination of RNA and protein structure during interaction

The complex between Rf protein and *orf147* transcript was modeled using homology modeling. The template used in the present modeling exercise was found with the help of BLAST (Basic Local Alignment Search Tool – (https://blast.ncbi.nlm.nih.gov/Blast.cgi) (Altschul *et al*., 1997). The sequence of the Rf protein (594 aa) protein was submitted as a query to the BLAST server, and the reference database contained entries from the PDB (RCSB Protein Data Bank) (Berman *et al*., 2000). The model was built with the help of Modeller version 9.16 (Šali and Blundell, 1993), and the superimposition figure of the template and model was generated using UCSF Chimera version 1.13.1 (Petterson *et al*., 2004).

### Functional validation of *Rf-PPR* in *Arabidopsis thaliana*

The putative Rfs determined after Y3H studies were further taken for functional validation in a model system *Arabidopsis thaliana.* The full-length coding sequence of the interacting *Rf* was cloned into pCAMBIA 2300 harboring GFP, NOS terminator and driven by AtAP_3_ promoter to generate a GFP fusion product (Supplementary Fig S3, Supplementary Table S4) and mobilized in *Agrobacterium tumefaciens* strain C58.

The T_1_ generation plants of previously developed transgenic *Arabidopsis thaliana* overexpressing *orf147* (OE-orf147)(Bhatnagar-Mathur *et al*., 2018) were transformed again using floral dip method (Zhang *et al*., 2006) and their seeds (hereafter referred as OE-orf147-Rf) collected after maturity. The collected seeds were then grown on appropriate media containing Ampicillin resistance for the selection of positives along with wild type (Col-0) and transgenic controls (OE-orf147). The surviving plantlets were transferred to autoclaved soil (1:1:1:1 ratio of black soil, red soil, compost and cocopeat) and grown under containment at 20°C, 16h day and 8h dark photoperiod and 65-70% RH. The genomic DNA from T_1_ generation plants overexpressing both *orf*-*Rf* genes (OE-orf147-Rf) was isolated using a NucleoSpin Plant II DNA isolation kit (Machery-Nagel, Germany) and screened for positives using Rf3-specific primers (Supplementary Table S2, Supplementary Table S3). The positive plants were then further observed for the number of silique formations compared to the WT plants.

### Subcellular localisation studies

The anthers from transgenic *Arabidopsis* plants from double-transformed (OE-orf147-Rf) and *orf147* expressing ones (OE-orf147) were obtained by gently tweaking the flower with forceps. The anther was stained with Mitotracker deep red (100nM) for 3 mins, followed by washing 5 times with distilled water. The anthers were subsequently fixed and mounted on the slide using 70% glycerol. GFP expression and co-localisation of the fusion product into the mitochondria were imaged using fluorescence (M16 FC, Leica) and confocal microscopy (TCS SP8, Leica) with Argon 488 nm and HeNe 633 nm laser wavelength for excitation of GFP and MitoTracker Deep Red.

### Expression profile analysis of positive *Arabidopsis* transgenics

RNA isolation of young flowers of *Arabidopsis* plants expressing OE-orf147-Rf, orf147 expressing (OE-orf147) and wild type Col-0 was carried out using the above-mentioned method. qRT-PCR analysis was carried out using gene-specific primers against *Rf3(*PP13) and *Orf147* (Supplementary Table S1, S2 and S3) using Realplex Real-Time PCR system (Eppendorf, Germany) with SYBR Green Real-Time PCR Master mix (Bioline) in 96 well optical reaction plates (Axygen, USA) sealed with ultra-clear sealing film (Platemax) and normalized using *Arabidopsis* reference primers (Supplementary Table S1). The relative expression levels of the genes were calculated using the 2^-ΔΔCt^ method, and the data is expressed as the mean ±SD (Winer *et al*., 1999).

## Results

### *In silico* identification of restorer genes and their relative expression in restorer and non-restorer lines

To identify the mitochondrial targeting genes, a total of 133 pigeonpea PPRs were identified from the PPR database, and after the removal of duplicates, 112 PPRs were obtained. *In silico* analysis determined that 22 of these PPRs were targeted to mitochondria and were mostly located on chromosomes 6,8,10, and 11 (Fig 1 A).

**Fig 1.**
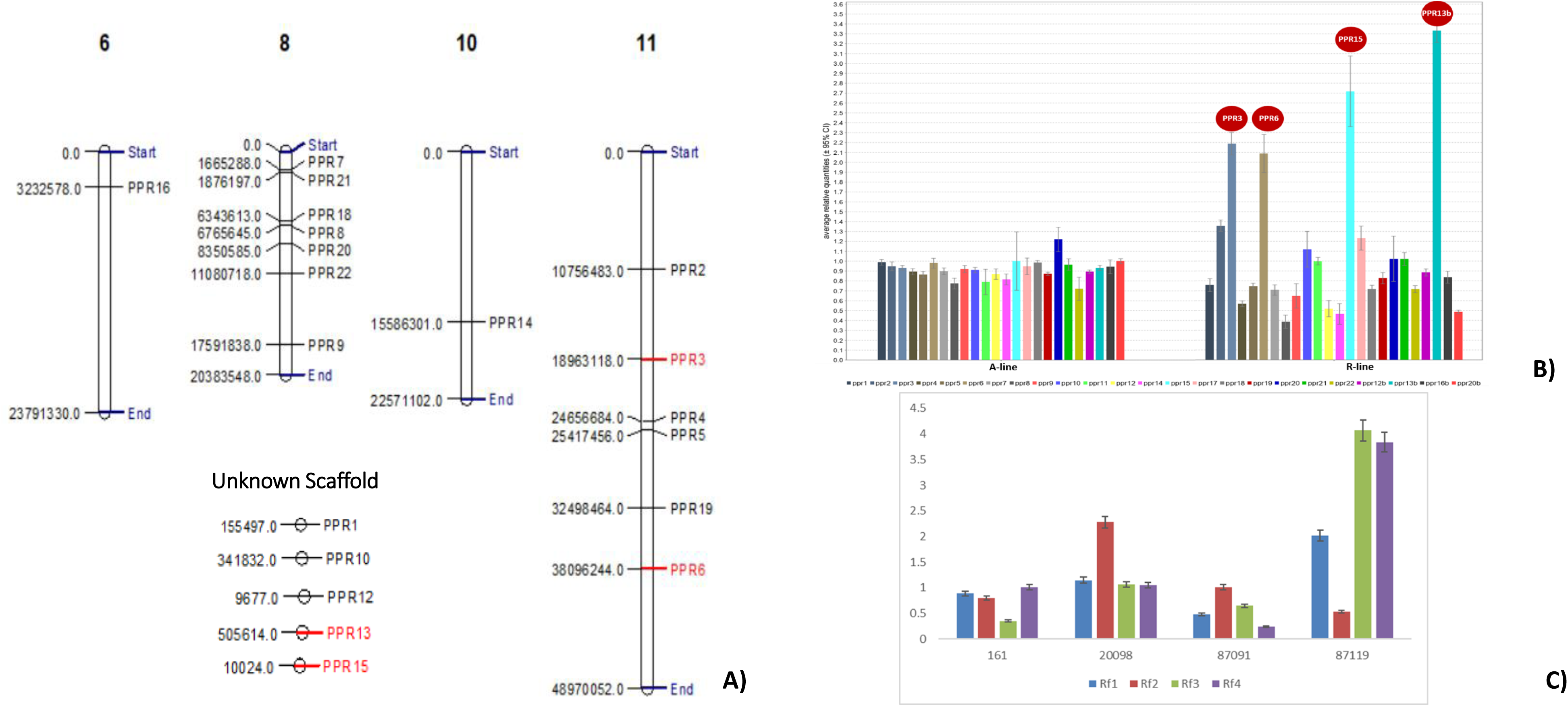
A) Chromosomal mapping of the identified PPRs. Most of the PPRs were found to be located in chromosome number 6, 8, 10 and 11. The four putative Rfs identified through expression profile analysis were observed to be located on chr no. 11 (Rf1 and Rf2) while the chromosomal location of Rf3 and Rf4 is not known. Image has been generated using MapChart and the chromosomal location has been obtained from NCBI corresponding to the PPR accessions. **B)** The identified 22 PPR genes were analysed for transcript expression level in the A-line (sterile line, ICPA 2039) and R-line (restorer line, ICPL 87119) of pigeonpea. Ideally, the possible Rfs would demonstrate lower transcript levels in the A-line while higher expression should be seen in the R-line. The genes showing maximum expression in R-line can be considered to be the putative restoration factors responsible for inducing restoration of fertility in pigeonpea. According to the expression profiles observed using qRT-PCR analysis, 4 PPRs have demonstrated highest level of expression in R-line while showing simultaneous low expression in A-line. These PPRs were selected and taken forward for interaction studies and validation**. C)** In addition to study of expression profiles in A and R-lines, the four PPRs selected were studied for transcript levels in other restorer and non-restorer lines as well. qRT-PCR analysis was performed with 4 genotypes. ICPL 87119 (Asha)-Restorer line, ICPL 161-Short duration restorer line, ICPL 20098-Medium duration restorer line and ICPL 87091-Non-restorer line. In this case, the 4 PPRs (Rf1, Rf2, Rf3 and Rf4) demonstrated low expression profile in the non-restorer line while exhibiting higher expression in restorer line further validating their role in restoration of fertility.

To confirm which ones of the 22 putative Rf-PPRs (Table 1) might potentially interact with *orf147* to inhibit its expression, identification of the ones that demonstrated upregulation in the restorer line (R-line) is crucial. Transcript profiling revealed expression of these genes at a lower level in the A-line (Fig 1B) compared to their expression in the restorer counterpart. Out of the 22 PPRs, 4 PPRS, *viz*, PPR3 (*Rf1*), PPR6 (*Rf2*), PPR13b (*Rf3*) and PPR15 (*Rf4*) showed significantly higher level of expression in R-line (Fig 1B). These selected PPRs were considered as potential Rfs and used for further studies.

**Table 1:**
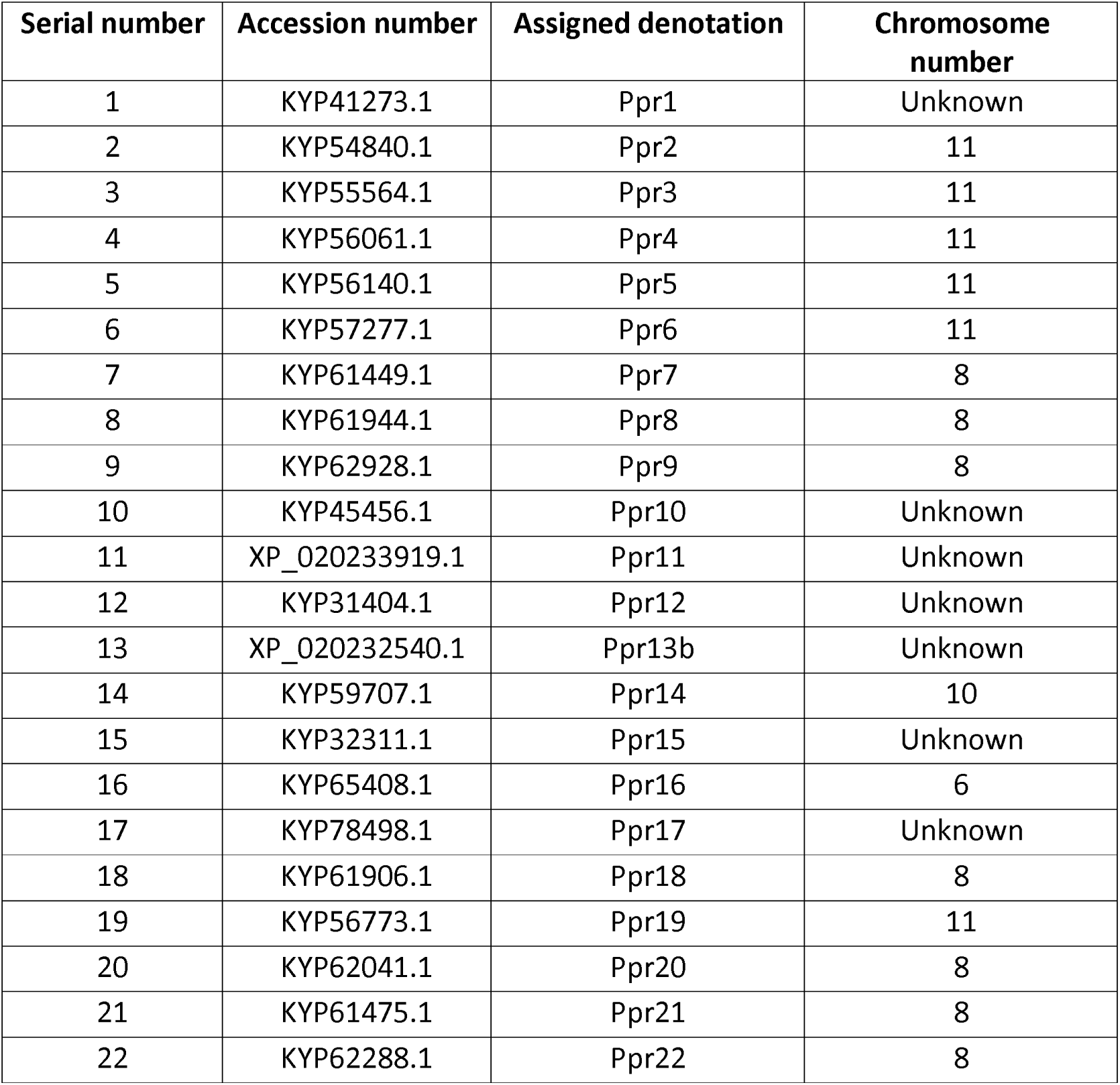
Sub-cellular localisation studies revealed 22 mitochondrial-targeting PPRs. The sub-cellular localisations of the 112 PPRs in mitochondria were identified through online software tools such as Mitoprot and Predotar tools. 22 were found to be mitochondrial targeting and were taken forward for expression analysis

In addition, comparison of expression profiles of *Rf1, Rf2, Rf3,* and Rf4 against other short-medium duration restorer and non-restorer lines of pigeonpea demonstrated the highest transcript levels in the corresponding restorer parent, ICPL 87119, and the least in the non-restorer line ICPL 87091. Short-duration restorer (ICPL 161) and medium-duration (ICPL 20098) restorer lines while captured higher transcripts than ICPL 87091; these were lower than ICPL 87119 (Fig 1C).

### Interaction between *Orf147*-*Rfs*

Based on expression analysis, 4 PPRs were chosen for interacting studies with *orf147*. Y3H study determined the appearance of blue colonies with Rf1, Rf3 and Rf4, indicative of a positive interaction between the RNA and protein components (Fig 2A). The empty fish vector control (AD) conferred cell growth on the DDO/X/A media, and the expression of the reporter gene was also detectable (Fig 2A). To rule out false positives, sequence analysis of the rescued plasmids with three *Rfs* was carried out that only confirmed *Rf3* as positive for Y3H (Supplementary Fig S4).

**Fig 2A:**
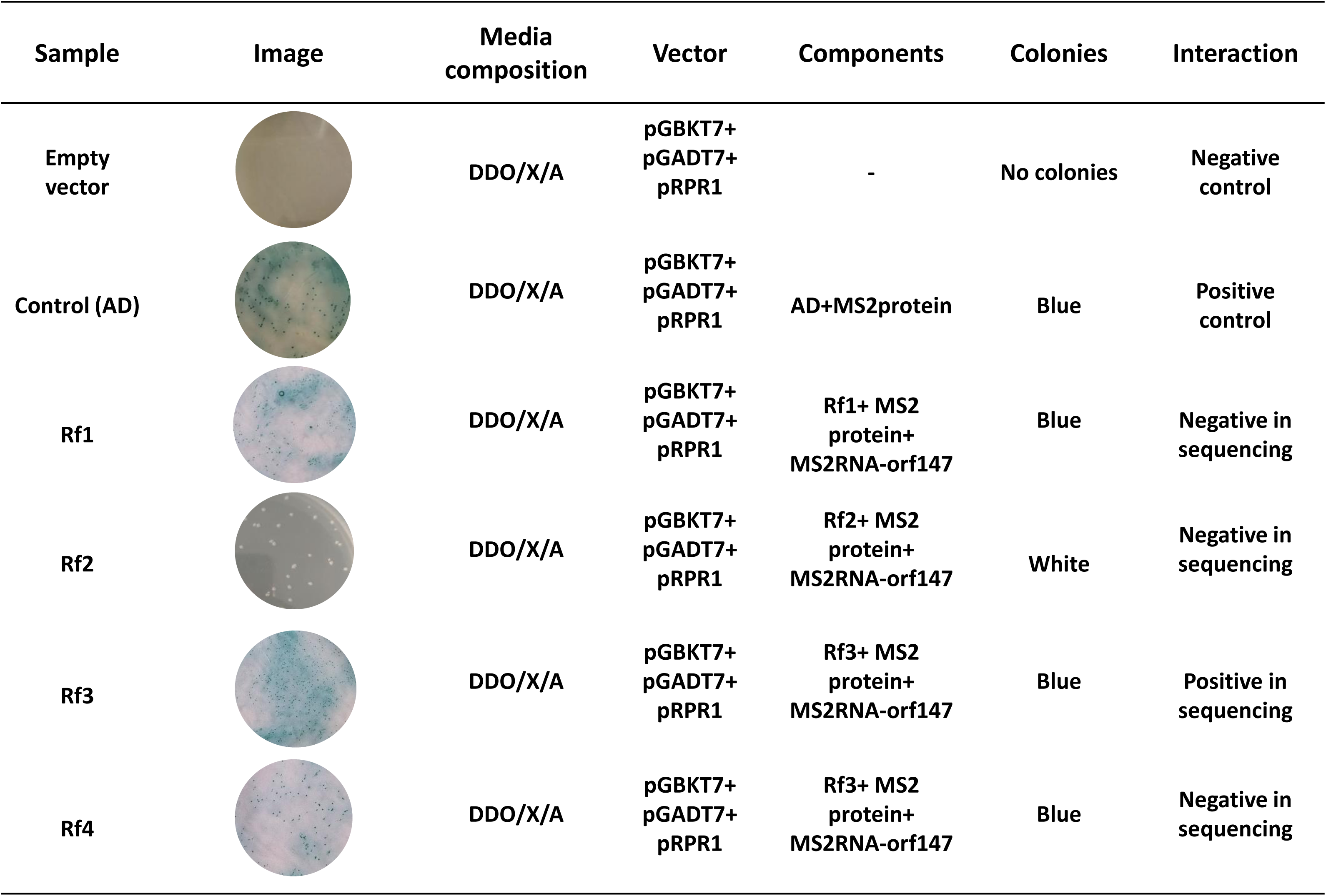
Interaction between *Orf147* and putative Rfs using yeast three hybrid studies. Yeast was also transformed with an empty vector and control AD. Co-transformation carried out with Rfs expressing in pGADT7 demonstrated occurrence of blue colonies in Rf1, Rf3 and Rf4. Sequence analysis demonstrated positive interaction of Rf3 with *Orf147*

### Homology modelling for RNA-protein interactions

The best result returned by the BLAST server contained an alignment between the query RF3 protein and chain A of PDB ID – 5I9D (Shen *et al*., 2016). This alignment was 434 amino acids long, having E-value = 4e-103, sequence identity = 38 % and sequence similarity = 59 %. PDB ID – 5I9D contains the crystal structure of a protein-RNA complex whose chain A is a pentatricopeptide repeat protein called dPPR-U8A2, and chain B is an RNA called U8A2. We built a homology model of the RF3 protein using the structure of dPPR-U8A2 as a template. The model built using Modeller version 9.16 (Šali and Blundell, 1993) omitted 75 amino acids from the N-terminus and 69 amino acids from the C-terminus of the RF3 protein because these residues were not covered by the template. The superimposition of the template and model is shown in Fig 2B. Rf protein, as well as dPPR-U8A2, bind a “UUAAUUU” motif in their substrate RNA that occurs in U8A2 as well as the *orf147* transcript. The “UUAAUUU” motif is located at position 413 – 419 in the sequence of the *orf147* transcript.

**Fig 2B:**
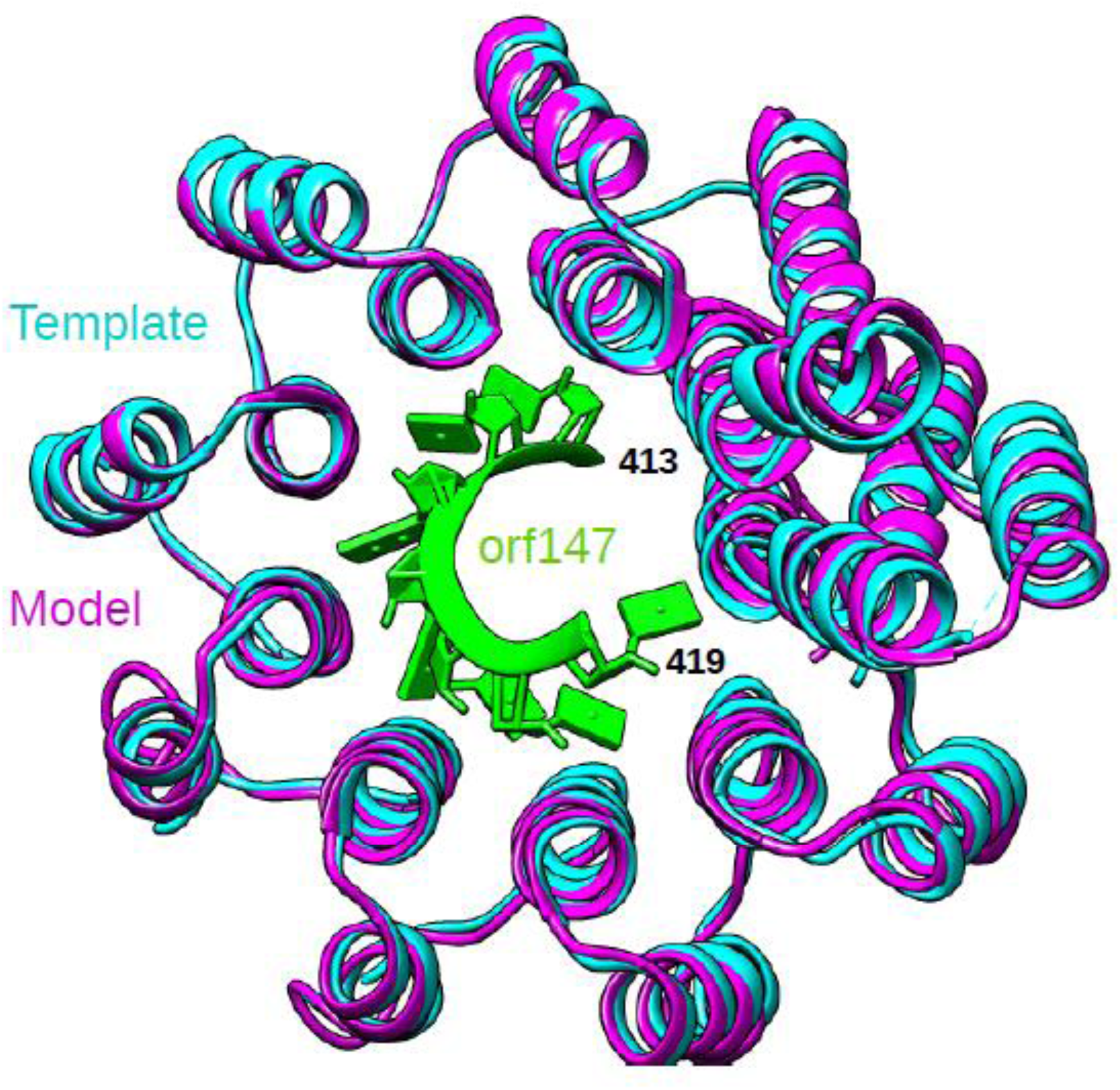
Homology modelling to demonstrate interaction between *Rf* and *orf147.* Superimposition of the model of the Rf protein (magenta) on the template dPPR-U8A2 (cyan). The UUAAUUU motif located at position 413 – 419 in the sequence of orf147 transcript (green) can also be seen. For the sake of clarity, loops occurring between amino acids 76 – 105, 176 – 212 and 503 – 525 in the model of Rf protein have been hidden.

### Functional validation of pigeonpea *Rf3* gene

The *Rf3* was functionally validated in *A. thaliana* model system by re-transforming the previously available transgenic *Arabidopsis* plants overexpressing pigeonpea *orf147* gene with *Rf3* driven by *AtAP3* promoter using the *Agrobacterium*-floral dip method.

Over 23 primary transgenic *Arabidopsis* events were generated expressing all three genes (*orf147* with the Rf3 gene and GFP reporter gene), out of which 14 events were confirmed to be PCR positive for both genes in the T_1_ generation (Supplementary Fig S5). The vegetative growth and morphology of the transgenic events were uniform and resembled with the wild types (Fig 3). While the *orf147* overexpressing transgenic events (OE-orf147) served as controls and were completely sterile, the number of siliques in doubly transformed events (OE-orf147-Rf) were comparable to the wild type plant (Table 2), implying restoration of fertility.

**Fig 3:**
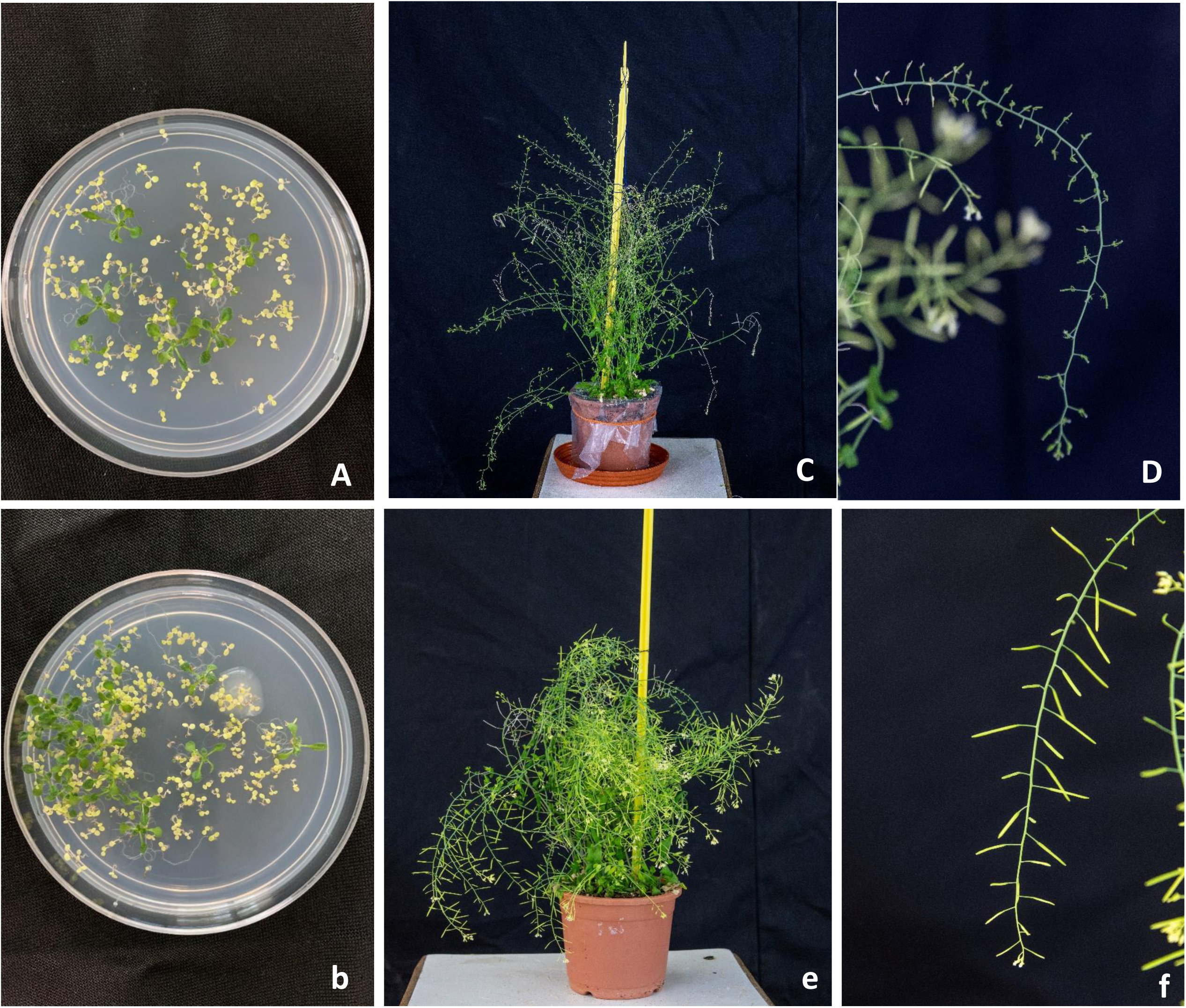
Showing sterile and fertility restored *A. thaliana* transgenics. *Orf147* transgenic *Arabidopsis thaliana* were double-transformed using *Rf3* construct by Floral dip method. The seeds were collected and selected on appropriate media with antibiotics (Ampicilin). The untransformed plants bleach out in selection media while the successful transformants demonstrated healthy growth. The healthy plants are selected and taken for further PCR analysis for identification of positive transformants. **A)** seeds of *Orf147* transgenic and **B)** double-transformed *Rf3* transgenic, placed on selection media **C)** and **D**) Control (*Orf147* transgenic) plants demonstrating sterility with absence of silique formation in T_2_ generation **E)** and **F**) Double-transformed *Rf3* transgenic plants showing healthy, silique formation in T_2_ generation. The development of siliques implies towards restoration of fertility

**Table 2:** Number of siliques in double transformed *Arabidopsis* transgenic and wild type. The number of siliques in the fertility-restored, double-transformed *A. thaliana* transgenics was found to be comparable to that of the wild-type.

| S.No. | Plant No | Number of siliques |
| --- | --- | --- |
| 1 | Wild type | 311 |
| 2 | #3 | 233 |
| 3 | #5 | 256 |
| 4 | #6 | 253 |
| 5 | #7 | 234 |
| 6 | #11 | 193 |
| 7 | #12 | 207 |
| 8 | #14 | 189 |
| 9 | #15 | 197 |
| 10 | #16 | 243 |
| 11 | #19 | 283 |
| 12 | #20 | 174 |
| 13 | #21 | 157 |
| 14 | #22 | 181 |
| 15 | #23 | 221 |

Quantitative real-time PCR (qRT-PCR) revealed that the expression level of *orf147* in the transgenic *Arabidopsis* (OE-orf147) was significantly higher (FC=3.4) than the double-transformed counterpart, OE-orf147-Rf. On the other hand, the latter showed a very high expression of Rf3 (FC=31), implying its role in the reduction in transcripts of *orf147* (Fig 4).

**Fig 4:**
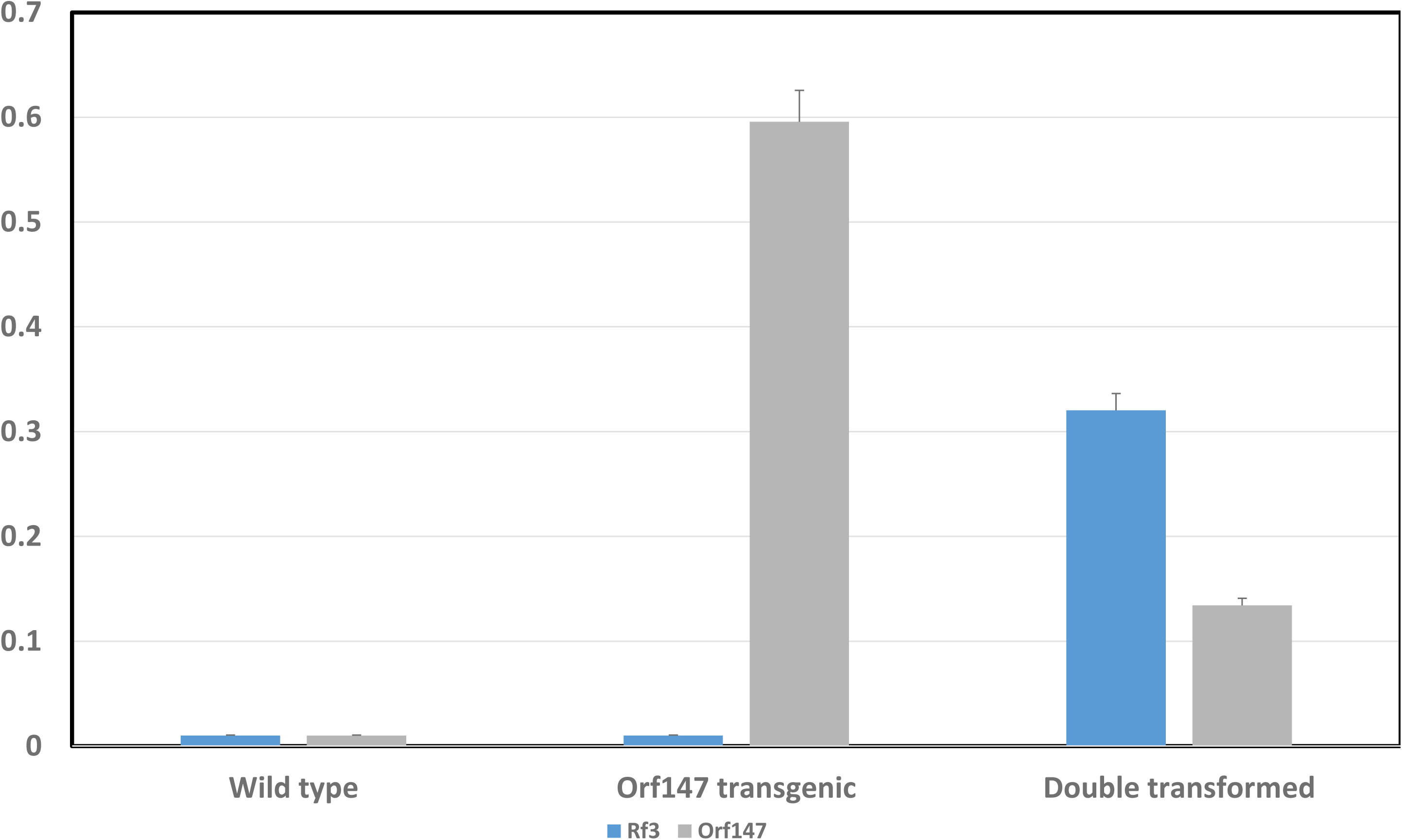
Expression profile of wild type, double transformed and *orf147* transgenic *Arabidopsis thaliana*. qRTPCR analysis demonstrated diminished expression of *orf147* in the double transformed *A. thaliana* and high expression of *Rf3.* This decrease in expression is attributed to the presence of *Rf3* which hinders the expression of *orf147* and eventually translation of cytotoxic protein.

### Sub-cellular localisation studies using fluorescence and confocal microscopy

Sub-cellular localisation studies were carried out to determine the localisation of RF3 in the cell. Preliminary studies using fluorescence microscopy showed successful incorporation of *Rf3* fused with GFP into the cell. Fluorescence was observed in the anthers of the double-transformed transgenic *Arabidopsis* (OE-orf147-Rf) plants (Fig 5). Subsequently, confocal microscopic studies of demonstrated green, fluorescent signal at 488nm and the same cell showed red fluorescent signal at 580nm (stained by MitoTracker Deep Red) in double transformed lines confirming the localisation of *Rf3* in the mitochondria (Fig 6 A-D).

**Fig 5:**
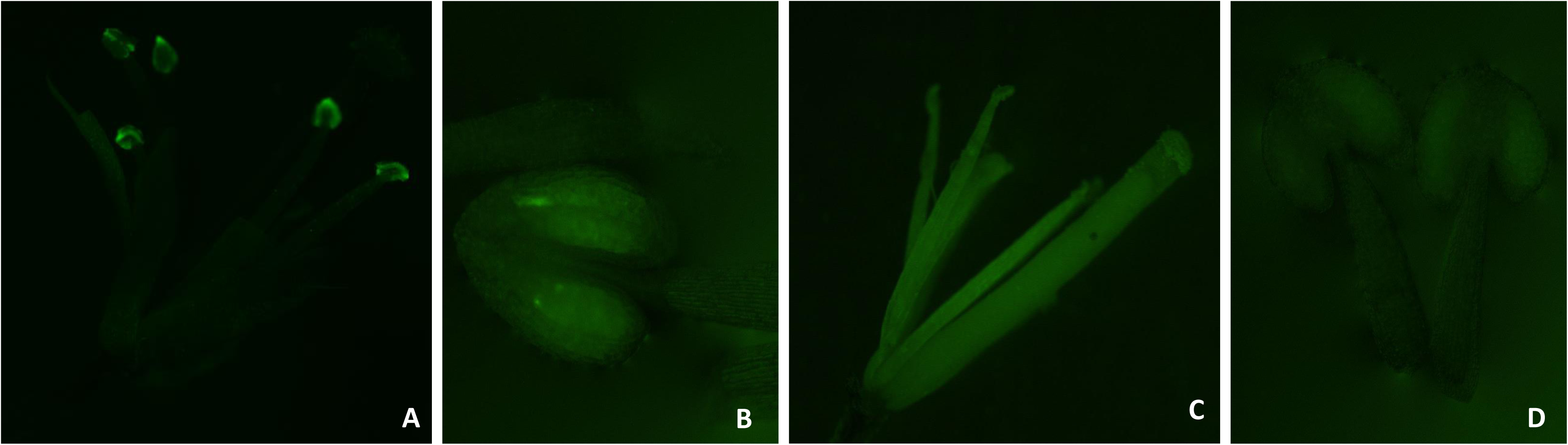
Fluorescence microscopy of double-transformed *Rf3* transgenics and control *orf147* transgenics. Due to the presence of Green fluorescent protein (GFP), fluorescence of observed in positive transformants and absence of fluorescence is seen in *orf147* transgenics. The flowers taken from both the samples were at similar stages to ensure uniformity. The study is a preliminary analysis of successful transformation. **A)** Fluorescence of double-transformed Rf3 transgenics and **B)** Fluorescence in single anther of Rf3 transgenic **C)** *Orf147* transgenic anther used as control shows no sign of fluorescence **D)** A closer view of the anther however shows faint traces of fluorescence which might be a result of autofluorescence.

**Fig 6:**
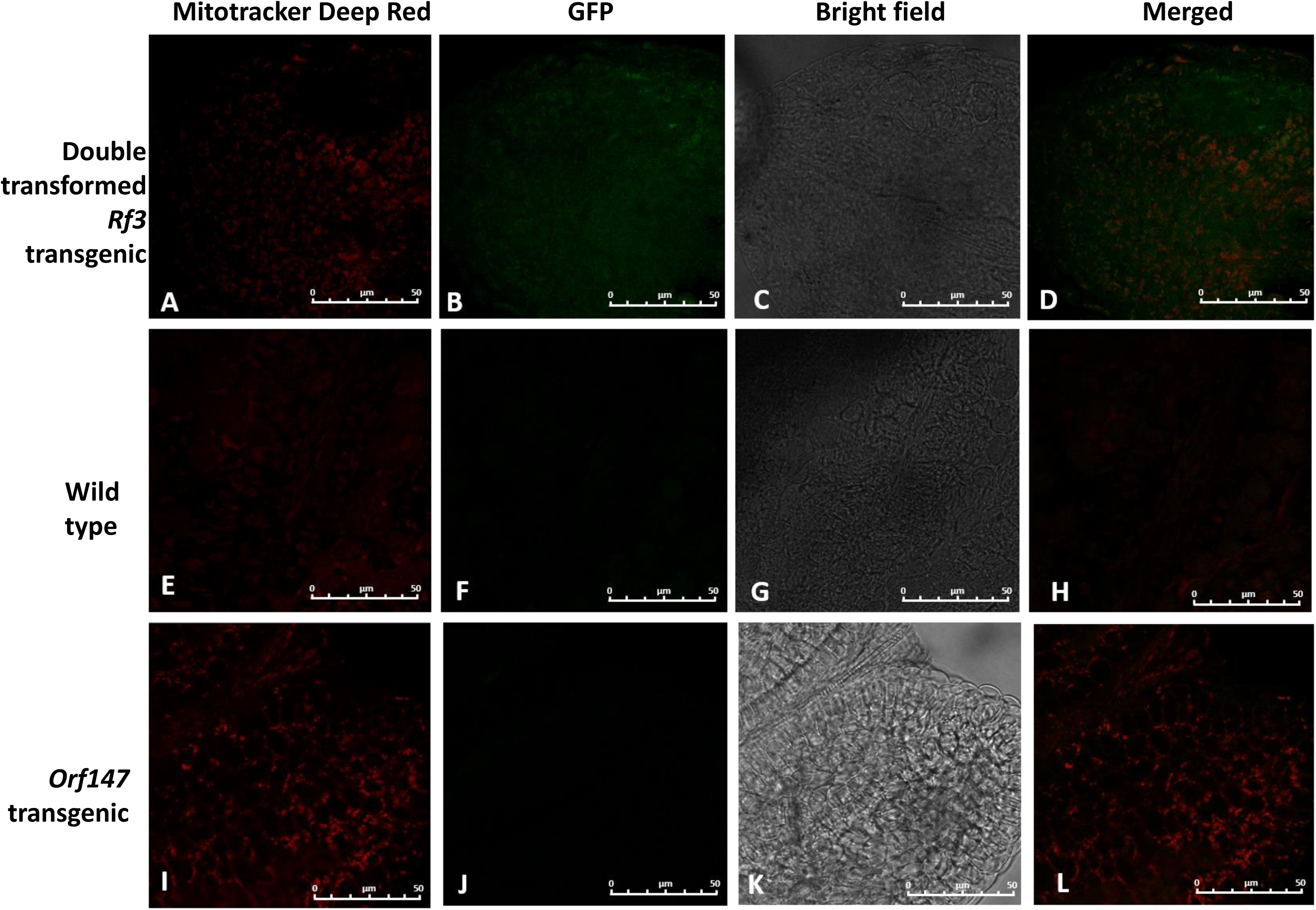
Confocal microscopy to study sub-cellular localisation. Double transformed Rf3 transgenic {(A-D), **A**: Mitotracker Deep Red, **B:** GFP, **C**: Bright Field, **D**: Merged}, Wild type {(E-H), **E**: Mitotracker Deep Red, **F**: GFP, **G**: Bright Field, **H**: Merged} and *orf147* transgenic {(I-L), **I**: Mitotracker Deep Red, **J**: GFP, **K**: Bright Field, **L**: Merged}

## Discussion

Pigeonpea improvement using breeding techniques has led to developments of over 140 varieties developed in the last 50 years across varying agroecological regions in India (Bohra *et al*., 2017). However, despite the evolving technologies, the average yield of pigeonpea inbreds remained below its potential yield of 2.5t/ha (Varshney *et al.,* 2012). To overcome these bottlenecks, the development of CMS-based hybrid systems were thrusted upon for pigeonpea improvement over the last few decades. The CMS lines derived from the A4 cytoplasm of *Cajanus cajanifolius* and the A2 cytoplasm of *Cajanus scarabaeoides* while have been extensively used to develop commercial hybrids, the success has been limited owing to the partial restoration of fertility and environment-genotype interactions (Choudhary and Singh, 2015). To mitigate these challenges, understanding the genetic basis of CMS and restoration of fertility, along with determining various factors responsible for fertility restoration, is critical for developing effective hybrid breeding systems in pigeonpea (Melonek *et al*., 2021).

This study is based on our previous work, where we had reported the CMS causing mitochondrial chimeric open reading frame (ORFs), *orf147* gene, in the A_4_ cytoplasm of *C. cajanifolius* (Bhatnagar-Mathur *et al*., 2018). Restorer of fertility (Rf) genes are nuclear genes which minimize the effect of CMS genes usually through post-transcriptional regulation of the CMS transcript, which subsequently reduces the CMS-inducing protein (Jaqueth *et al*., 2020; Dahan and Mireau, 2013). Using genome-wide analyses of the pigeonpea PPR family, we narrowed down the interacting partners with the CMS-causing *orf147* in pigeonpea. *In silico* studies performed in plant species have demonstrated that most of the Rfs identified belong to a sub-group of P-type PPRs *viz* restorer of fertility-like PPR (RFL) proteins (Dahan and Mireau, 2013; Sykes *et al*., 2017; Barchenger *et al*., 2018; Kubo *et al*., 2020). A total of 112 PPRs of pigeonpea were identified from the pigeonpea PPR database. Since Rfs are mitochondrial targeting, the sub-cellular localisation of the 112 PPRs was analysed using online tools like Predotar, and Mitoprot, which shortlisted 22 PPRs as potential Rfs. Chromosomal mapping of the selected Rfs indicated most of these residing in chromosome numbers 6,8,10, and 11, while few belonged to unknown scaffolds.

Furthermore, using the three-hybrid system of pigeonpea consisting of sterile A-line (ICPA 2039), maintainer B-line (ICPB 2039) and restorer R-line (ICPL 87119), the expression profiles of 22 PPR genes were studied in A-line and R-lines. Expression analyses in flower buds of pigeonpea indicated similar expression levels of most PPRs in both A-line and R-line with the exception of 4 (PPR3, PPR6, PPR13 and PPR15) that showed contrasting expression profiles. This did not come as a surprise as Rfs, being restoration factors, have higher expression in the R-line compared to the A-line. Owing to these differential expression levels, the selected four PPRs were further evaluated for their restoration capabilities. The expression levels of these 4 PPRs were confirmed in several other restorer and non-restorer lines of pigeonpea; that revealed the least expression in the non-restorer line (ICPL 87091), intermediate levels in medium duration restorer line (ICPL 20098), and showed highest expression in the long-duration restorer line ICPL 87119 implying towards their possible role as restoration factors in a range of restorer backgrounds.

Studies have established the RNA-binding properties of PPRs (Barkan and Small, 2014) due to the presence of side chains lining the central groove that renders them hydrophilic, thereby suggesting that PPRs are RNA-binding rather than protein-binding (Bentolila *et al*., 2002). To confirm the binding specificity of these potential restorer factors to *orf147*, we adapted the yeast-three hybrid system studies by generating three components where along with a bait (MS2 protein in pGBKT7) and prey (individual PPRs in pGADT7) vector, a third hybrid ligand (MS2RNA-orf147 in pRPR1) was involved. Out of the 4 PPRs, PPR13 (Rf3) showed positive sequencing results and was therefore deemed the “putative restoration factor” against *orf147*.

The exact binding site of *orf147* to PPR13 was predicted by homology modelling, which helped create the Rf-orf147 complex. The complex showed binding of Rf protein to the substrate *orf147* RNA at a “UUAAUUU” motif located at positions 413-419 of the RNA substrate. Predictions of PPR motif folding demonstrated that these fold into super helical ribbon-like sheets with the C terminal of the helix constituting the external surface of protein while the N terminal forms the RNA-binding side (Barkan and Small, 2014). These results were significant as they validate the RNA-binding characteristic of the Rf to *orf147* and the exact location, along with providing a basis for additional analysis. The first instance of mitochondrial targeting and binding of PPRs was revealed in the Rf-PPR592 of petunia, which actively reduces the CMS-causing PCF protein by binding with the *pcf* transcript (Gillman *et al*., 2007). The binding characteristic of Rfs has also been reported previously in BT-type cytoplasmic male sterile rice (*Oryza sativa* L.) wherein the restorer protein, Rf1, binds to CMS causing *orf79* transcript and restoration of fertility occurs via RNA processing thereafter (Kazama *et al*., 2008). RNA editing and cleavage of the *atp6-orf79* has also been observed by Rf1A and Rf1B in rice (Wang *et al*., 2006). In a related study, another restorer protein, Rf5, corresponding to *atp6-orf79,* was identified in HL-rice wherein the role of an accessory protein, GRP162, in RNA binding and subsequent cleavage of the transcript, was established (Hu *et al*., 2012).

Subsequently, putative restoration factor (PPR13; Rf3) was functionally validated using transgenic expression studies, where T_1_ generation plants of double transformed transgenic *Arabidopsis thaliana* plants that overexpressed *orf147* and AtAP3:Rf3: GFP-Nos. While the overexpression of *orf147* alone exhibited partial sterility in T_1_ lines, and complete sterility in T_2_ generation (Bhatnagar-Mathur *et al*., 2018), the plants that co-expressed both *orf147* and Rf3 produced siliques in numbers comparable to the wild types. This study indicated the restoration of fertility in the otherwise sterile *A. thaliana* plants, validating the restoration capacity of *Rf3*. Moreover, the positive double-transformed *A. thaliana* transgenics events (OE-orf147-Rf) demonstrated phenology similar to their WT counterparts. Interestingly, qRT-PCR studies revealed reduced levels of *orf147* transcripts in the double-transformed *Rf3* line, while expression levels of *Rf3* were significantly higher. This implied suppressed activity of the *orf147* due to the presence of Rf3 and, hence, restoration of fertility.

Our study further elucidated the mechanism associated with the observed restoration of fertility in *Arabidopsis* plants, as well as sub-cellular localisation analysis using a fused GFP-Rf3 construct. Fluorescence microscopy demonstrated that the gene was targeting the flower’s anther, and the gene’s localization in the mitochondria was further confirmed using confocal studies. This significant result confirmed the inherent mitochondrial targeting ability of restoration factors. The Rf proteins bind specifically to CMS transcripts in the mitochondria, thereby reducing the production of CMS-inducing proteins (Gaborieau *et al*., 2016)

Based on these results, our study for the first time report *Rf3* as the corresponding restoration factor against *orf147* in pigeonpea. Further studies in the native pigeonpea system by introducing Rf3 in A-line would also validate the function of the Rf3 protein in the crop system. This is of utmost significance as it can be translated into the generation of stable hybrids in this important “opportunity” legume crop.

### Concluding comments

The identification of a restorer gene associated with fertility restoration in the A4 CMS system in pigeonpea and the proof of interaction between CMS-causing orf147 and Rf3 deduce the underlying mechanism, opening a wider avenue for further studies in other legumes to establish robust inducible hybrid systems.

## Abbreviations

CMS: Cytoplasmic male sterility
Col-0: Columbia-0
DDO/X/A: Double dropout media / X-α-gal/ Aureobasidin A
GFP: Green fluorescent protein
qRT-PCR: quantitative real-time PCR
PPR: Pentatricopeptide repeat protein
Rf: Restoration factors
WT: Wild type
Y3H: Yeast three hybrid system

## Acknowledgments

The authors thank Dr Anupama Hingane (former Scientist, Pigeonpea Breeding Unit, International Crops Research Institute for the Semi-Arid Tropics, India) for providing the seeds of restorer lines and Mr Jismon Jose (Indian Institutes of Science Education and Research, Tirupati) for assistance with confocal microscopy.

## Author contribution

**JB:** Formal analysis, methodology, data curation, investigation, writing-original draft, visualization; **RBN:** Data curation, visualization, methodology, review and editing; **DSR:** Data curation, conceptualization, methodology, review and editing; **VS:** Data curation, review and editing; **YR:** methodology; Formal analysis, Data curation; **RY:** Formal analysis; **YK:** Investigation and review; **PR**: Formal analysis **PBM**: Conceptualization, Funding, Project administration, review and funding acquisition; **PSR:** Conceptualization, Supervision, Project administration and review.

## Supplementary data

The following supplementary data are available at JXB online.

**Table S1:** List of primers of PPRs used for qRT-PCR studies.

**Table S2:** PCR program used for the present study.

**Table S3:** List of primers and gene sequences used for cloning of components of the Y3H study

**Table S4:** List of primers used for cloning AtAP3, GFP-Nos and Rf3 into pCAMBIA 2300

**Fig S1:** A) pRPR1 (1X TetO) _gRNA_handle_RPT1t expressing MS2RNA-Orf146 B) pGBKT7 expressing MS2 protein (7303bp) C) pGADT7 expressing Rfs

**Fig S2:** Cloning strategy of A) MS2 protein expressing in pGBKT7 bait vector and B) MS2orf147 hybrid RNA expressing in pRPR1 vector

**Fig S3:** Cloning strategy of Rf3 into pCAMBIA2300 harbouring GFP-NOS and driven by AtAP3 promoter

**Fig S4:** Sequence analysis was performed of the rescued plasmid against Rf3 and Orf147, which constitute two of the components of the Y3H study, using clustal omega, and the chromatogram quality was checked using Genious software. Sequence alignment of i) Rf3 and ii) orf147 and respective chromatograms showing distinct peaks

## References

Altschul SF, Madden TL, Schäffer AA, Zhang J, Zhang Z, Miller W, Lipman DJ. 1997. Gapped BLAST and PSI-BLAST: a new generation of protein database search programs. Nucleic acids research 25(17), 3389–402.

Barchenger DW, Said JI, Zhang Y, Song M, Ortega FA, Ha Y, Kang BC, Bosland PW. 2018. Genome-wide identification of Chile pepper pentatricopeptide repeat domains provides insight into fertility restoration. Journal of the American Society for Horticultural Science 143, 418–429.

Barkan A, Small I. 2014. Pentatricopeptide repeat proteins in plants. Annual Review of Plant Biology 65, 415–442.

Bentolila S, Alfonso AA, Hanson MR. 2002. A pentatricopeptide repeat-containing gene restores fertility to cytoplasmic male-sterile plants. Proceedings of the National Academy of Sciences of the United States of America 99, 10887–10892.

Berman HM, Westbrook J, Feng Z, Gilliland G, Bhat TN, Weissig H, Shindyalov IN, Bourne PE. 2000. The protein data bank. Nucleic acids research 28(1), 235–42.

Bhatnagar-Mathur P, Gupta R, Reddy PS, Reddy BP, Reddy DS, Sameer kumar C V., Saxena RK, Sharma KK. 2018. A novel mitochondrial *orf147* causes cytoplasmic male sterility in pigeonpea by modulating aberrant anther dehiscence. Plant Molecular Biology 97, 131–147.

Bhattacharya J, Reddy DS, Prasad K, Nitnavare RB, Bhatnagar-Mathur P, Sudhakar P. 2023. Ectopic expression of pigeonpea *Orf147* gene imparts partial sterility in *Cicer arietinum*. Gene 868, 147372.

Bohra A, Jha R, Pandey G, et al. 2017. New hypervariable SSR markers for diversity analysis, hybrid purity testing and trait mapping in pigeonpea [*Cajanus cajan* (L.) Millspaugh]. Frontiers in Plant Science 8, 1–15.

Brown SE, Ross MF, Sanjuan-Pla A, Manas AR, Smith RA, Murphy MP. 2007. Targeting lipoic acid to mitochondria: synthesis and characterization of a triphenylphosphonium-conjugated α-lipoyl derivative. Free Radical Biology and Medicine 42(12), 1766–80.

Chase CD. 2007. Cytoplasmic male sterility: a window to the world of plant mitochondrial-nuclear interactions. Trends in Genetics 23, 81–90.

Chen Z, Zhao N, Li S, Grover CE, Nie H, Wendel JF, Hua J. 2017. Plant mitochondrial genome evolution and cytoplasmic male sterility. Critical Reviews in Plant Sciences 36, 55–69.

Choudhary AK, Singh IP. 2015. A study on comparative fertility restoration in A_2_ and A_4_ cytoplasms and its implication in breeding hybrid pigeonpea [*Cajanus cajan* (L.) Millspaugh]. American Journal of Plant Sciences 06, 385–391.

Dahan J, Mireau H. 2013. The Rf and Rf-like PPR in higher plants, a fast-evolving subclass of PPR genes. RNA Biology 10, 1286–1293.

Dalvi VA, Saxena KB, Luo RH, Li YR. 2010. An overview of male-sterility systems in pigeonpea [*Cajanus cajan* (L.) Millsp.]. Euphytica 173, 397–407.

Desloire S, Gherbi H, Laloui W, et al. 2003. Identification of the fertility restoration locus, *Rfo*, in radish, as a member of the pentatricopeptide-repeat protein family. EMBO reports 4(6), 588–94.

FAO. 2021. International Production : Pigeon Peas. https://agriexchange.apeda.gov.in/International_Productions/International_Production.aspx?ProductCode=0197#:~:text=International%20production%20of%20Pigeon%20Peas,All. Accessed April, 2024.

Fatokimi EO, Tanimonure VA. 2021. Analysis of the current situation and future outlooks for pigeon pea (*Cajanus Cajan* ) production in Oyo State, Nigeria: A Markov Chain model approach. Journal of Agriculture and Food Research 6, 100218.

Gaborieau L, Brown GG, Mireau H. 2016. The propensity of pentatricopeptide repeat genes to evolve into restorers of cytoplasmic male sterility. Frontiers in Plant Science 7, 1–10.

Gillman JD, Bentolila S, Hanson MR. 2007. The petunia restorer of fertility protein is part of a large mitochondrial complex that interacts with transcripts of the CMS-associated locus. Plant Journal 49, 217–227.

Hamid NAA, Ismail I. 2018. PEG-4000 increased the mating efficiency of yeast-two hybrid screening process using PmF-box1 as Bait. Sains Malaysiana 47, 2961–2968.

Hu J, Wang K, Huang W, et al. 2012. The rice pentatricopeptide repeat protein RF5 restores fertility in Hong-Lian cytoplasmic male-sterile lines via a complex with the glycine-rich protein GRP162. Plant Cell 24, 109–122.

Jaqueth JS, Hou Z, Zheng P, Ren R, Nagel BA, Cutter G, Niu X, Vollbrecht E, Greene TW, Kumpatla SP. 2020. Fertility restoration of maize CMS-C altered by a single amino acid substitution within the *Rf4 bHLH* transcription factor. Plant Journal 101, 101–111.

Kazama T, Nakamura T, Watanabe M, Sugita M, Toriyama K. 2008. Suppression mechanism of mitochondrial ORF79 accumulation by Rf1 protein in BT-type cytoplasmic male sterile rice. Plant Journal 55, 619–628.

Klein RR, Klein PE, Mullet JE, Minx PA, Rooney WL, Schertz KF. 2005. Fertility restorer locus *Rf1* of sorghum (*Sorghum bicolor* L.) encodes a pentatricopeptide repeat protein not present in the colinear region of rice chromosome 12. Theoretical and Applied Genetics 111, 994–1012.

Koizuka N, Imai R, Fujimoto H, Hayakawa T, Kimura Y, Kohno-Murase J, Sakai T, Kawasaki S, Imamura J. 2003. Genetic characterization of a pentatricopeptide repeat protein gene, *orf687*, that restores fertility in the cytoplasmic male-sterile Kosena radish. The Plant Journal 34(4), 407–15.

Kubo T, Arakawa T, Honma Y, Kitazaki K. 2020. What does the molecular genetics of different types of restorer-of-fertility genes imply?. Plants 13, 9(3):361.

Melonek J, Duarte J, Martin J, et al. 2021. The genetic basis of cytoplasmic male sterility and fertility restoration in wheat. Nature Communications 12.

Pettersen EF, Goddard TD, Huang CC, Couch GS, Greenblatt DM, Meng EC, Ferrin TE. 2004. UCSF Chimera—a visualization system for exploratory research and analysis. Journal of computational chemistry 25(13),1605–12.

Šali A, Blundell TL. 1993. Comparative protein modelling by satisfaction of spatial restraints. Journal of molecular biology 234(3),779–815.

Saxena KB, Kumar R V., Srivastava N, Shiying B. 2005. A cytoplasmic-nuclear male-sterility system derived from a cross between *Cajanus cajanifolius* and *Cajanus cajan*. Euphytica 145, 289–294.

Saxena KB, Kumar RV, Tikle AN, et al. 2013. ICPH 2671 - the world’s first commercial food legume hybrid. Plant Breeding 132, 479–485.

Saxena KB, Sharma D, Vales MI. 2017. Development and commercialization of CMS pigeonpea hybrids. Plant Breeding Reviews 41, 103–167.

Shen C, Zhang D, Guan Z, Liu Y, Yang Z, Yang Y, Wang X, Wang Q, Zhang Q, Fan S, Zou T. 2016. Structural basis for specific single-stranded RNA recognition by designer pentatricopeptide repeat proteins. Nature communications 7(1), 11285.

Sinha P, Singh VK, Suryanarayana V, Krishnamurthy L, Saxena RK, Varshney RK. 2015 Evaluation and validation of housekeeping genes as reference for gene expression studies in pigeonpea (*Cajanus cajan*) under drought stress conditions. PloS one 10,4.

Sykes T, Yates S, Nagy I, Asp T, Small I, Studer B. 2017. In silico identification of candidate genes for fertility restoration in cytoplasmic male sterile perennial ryegrass (*Lolium perenne* L.). Genome Biology and Evolution 9, 351–362.

Varshney RK, Chen W, Li Y et al. 2012. Draft genome sequence of pigeonpea (*Cajanus cajan*), an orphan legume crop of resource-poor farmers. Nature biotechnology 30(1),83.

Wang Z, Zou Y, Li X, et al. 2006. Cytoplasmic male sterility of rice with Boro II cytoplasm is caused by a cytotoxic peptide and is restored by two related PPR motif genes via distinct modes of mRNA silencing. Plant Cell 18, 676–687.

Winer J, Jung CKS, Shackel I, Williams PM. 1999. Development and validation of real-time quantitative reverse transcriptase-polymerase chain reaction for monitoring gene expression in cardiac myocytes in vitro. Analytical Biochemistry 270, 41–49.

Zhang X, Henriques R, Lin SS, Niu QW, Chua NH. 2006. *Agrobacterium*-mediated transformation of *Arabidopsis thaliana* using the floral dip method. Nature Protocols 1, 641– 646.

